# Genetic determinants of receptor-binding preference and zoonotic potential of H9N2 avian influenza viruses

**DOI:** 10.1101/2020.05.11.088203

**Authors:** Thomas P. Peacock, Joshua E. Sealy, William T. Harvey, Donald J. Benton, Richard Reeve, Munir Iqbal

**Affiliations:** Department of Infectious Diseases, Imperial College London, UK, W2 1NY; Avian Influenza Group, The Pirbright Institute, Woking, UK, GU24 0NF; Royal Veterinary College, University of London, UK, NW1 0TU; Boyd Orr Centre for Population and Ecosystem Health, Institute of Biodiversity, Animal Health and Comparative Medicine, College of Medical, Veterinary and Life Sciences, University of Glasgow, UK, G12 8QQ; The Francis Crick Institute, London, UK, NW1 1ST

## Abstract

Receptor recognition and binding is the first step of viral infection and a key determinant of host specificity. The inability of avian influenza viruses to effectively bind human-like sialylated receptors is a major impediment to their efficient transmission in humans and pandemic capacity. Influenza H9N2 viruses are endemic in poultry across Asia and parts of Africa where they occasionally infect humans and are therefore considered viruses with zoonotic potential. We previously described H9N2 viruses, including several isolated from human zoonotic cases, showing a preference for human-like receptors. Here we take a mutagenesis approach, making viruses with single or multiple substitutions in H9 haemagglutinin to determine the genetic basis of preferences for alternative avian receptors and for human-like receptors. We describe amino acid motifs at positions 190, 226 and 227 that play a major role in determining receptor specificity, and several other residues such as 159, 188, 193, 196, 198 and 225 play a smaller role. Furthermore, we show changes at residues 135, 137, 147, 157, 158, 184, 188, and 192 can also modulate virus receptor avidity and that substitutions that increased or decreased the net positive charge around the haemagglutinin receptor-binding site show increases and decreases in avidity, respectively. The motifs we identify as increasing preference for the human-receptor will help guide future H9N2 surveillance efforts and facilitate our understanding of the emergence of influenza viruses with high zoonotic potential.

**Author Summary:** As of 2020, over 60 infections of humans by H9N2 influenza viruses have been recorded in countries were the virus is endemic. Avian-like cellular receptors are the primary target for these viruses. However, given that human infections have been detected on an almost monthly basis since 2015, there may be a capacity for H9N2 viruses to evolve and gain the ability to target human-like cellular receptors. Here we identify molecular signatures that can cause viruses to bind human-like receptors, and we identify the molecular basis for the distinctive preference for sulphated receptors displayed by the majority of recent H9N2 viruses. This work will help guide future surveillance by providing markers that signify the emergence of viruses with enhanced zoonotic potential as well as improving understanding the basis of influenza virus receptor-binding.

## Introduction

In the 1990s avian influenza virus subtypes H5N1 and H9N2 underwent a host-switch from wild birds to domestic poultry where they have circulated ever since. H9N2 has since become one of the most widespread strains in poultry, infecting domestic fowl throughout Asia and North Africa, where it circulates hyper-endemically [1–4]. Zoonotic H9N2 cases are also occasionally detected, with human infections reported in Hong Kong, mainland China, Bangladesh, Egypt, Pakistan, Oman, Indian and Senegal; over half of human infections have been reported in the last 4 years alone, all of which indicates a growing pandemic threat from these viruses [4–12]. Although no human-to-human transmission has been recorded, some H9N2 virus strains have shown a high propensity for airborne transmission between ferrets [13, 14], the most commonly used model for human influenza transmission.

The influenza glycoprotein, haemagglutinin (HA), mediates attachment of virus to host cells through binding of glycans with terminal sialic acids moieties. The human upper respiratory tract (URT) is rich in glycans with terminal α2,6-linked sialic acid (SA) and is the primary site of replication for human influenza viruses. An important determinant for adaptation to the human URT is the ability of HA to bind α2,6-linked sialylated glycans [15]. However, most avian influenza viruses preferentially bind to glycans with terminal α2,3-linked SAs, which are common in the avian gastrointestinal and respiratory tracts [16]. Therefore, for avian influenza viruses to be able to efficiently infect and transmit between humans they must gain the ability to bind to α2,6-linked SA.

Several molecular determinants have been shown to influence the receptor-binding profile of H9N2 viruses, including position 226 (H3 HA numbering used throughout; position 216 in mature polypeptide H9 numbering) – in one strain, Q226L alone could facilitate greater replication of an H9N2 isolate in human epithelial airway cells [17]. Furthermore, substitutions at positions 155, 190 and 227 (145, 180 and 217 in H9 numbering) have also been shown to play a role in receptor-binding preference in some H9N2 viruses [13, 18–20]. However, understanding of H9N2 receptor-binding preference remains piecemeal, with no studies having systematically looked at the roles of multiple residues, or residue combinations, in the variation in H9N2 receptor preference.

In a previous study, we tested the receptor-binding preference of several H9N2 viruses and described notable receptor preference variability amongst circulating H9N2 viruses which we hypothesised was due to amino acid variability at residues 190, 226 and 227 [21]. In this study we take three H9N2 viruses that are representative of different receptor-binding profiles, including a virus isolated from a human with a natural preference for α2,6-linked SA, and generate recombinant virus libraries with HA amino acid substitutions that represent reciprocal changes between the progenitor H9N2 viruses. We test the receptor-binding of these libraries and show residues 190, 226, 227, and to a lesser extent 159, 188, 193, 196, 198, and 225, explain this receptor preference variability. We further show several antibody escape mutants have changed receptor preference or avidity, and we describe a correlation between the electrostatic charge of the HA head and receptor avidity. Finally, we use the insights from these experiments to predict that certain H9N2 lineages may have a naturally high propensity to bind human receptors and therefore possess a higher zoonotic potential within the general viral population.

## Results

Three previously characterised viruses were chosen to act as mutagenesis backgrounds due to their distinct receptor-binding phenotypes. Receptor-binding profiles were measured using three receptor analogues: sulphated and non-sulphated 3’sialyllactosamine (3SLN(6su) and 3SLN, respectively) and 6’sialyllactosamine (6SLN). Both 3SLN(6su) and 3SLN are analogues for avian-like receptors while 6SLN is an analogue for human-like receptors. The virus A/chicken/Pakistan/UDL-01/2008 (UDL1/08) displays high binding avidity to 3SLN(6su), but not 3SLN (avian-like), with residual binding to the human-like receptor 6SLN (Figure 1A), similar to the majority of contemporary H9N2 viruses [20–22]. The virus A/chicken/Emirates/R66/2002 (Em/R66) binds to both 3SLN(6su) and 3SLN but has no detectable binding to 6SLN (Figure 1B), similar to conventional avian-adapted H5N1 and H7 viruses [21, 23]. Finally, A/Hong Kong/33982/2009 (HK/33982) binds to all three receptor analogues, but with an appreciable preference for 6SLN, similar to early human pandemic H3N2 viruses and zoonotic H7N9 viruses (Figure 1C) [21, 24]. To test the molecular basis of these different receptor preferences, libraries of individual or multiple reciprocal mutants were generated between these viruses with a particular focus on positions 190, 226 and 227, as well as several other nearby RBS residues.

**Figure 1 -.**
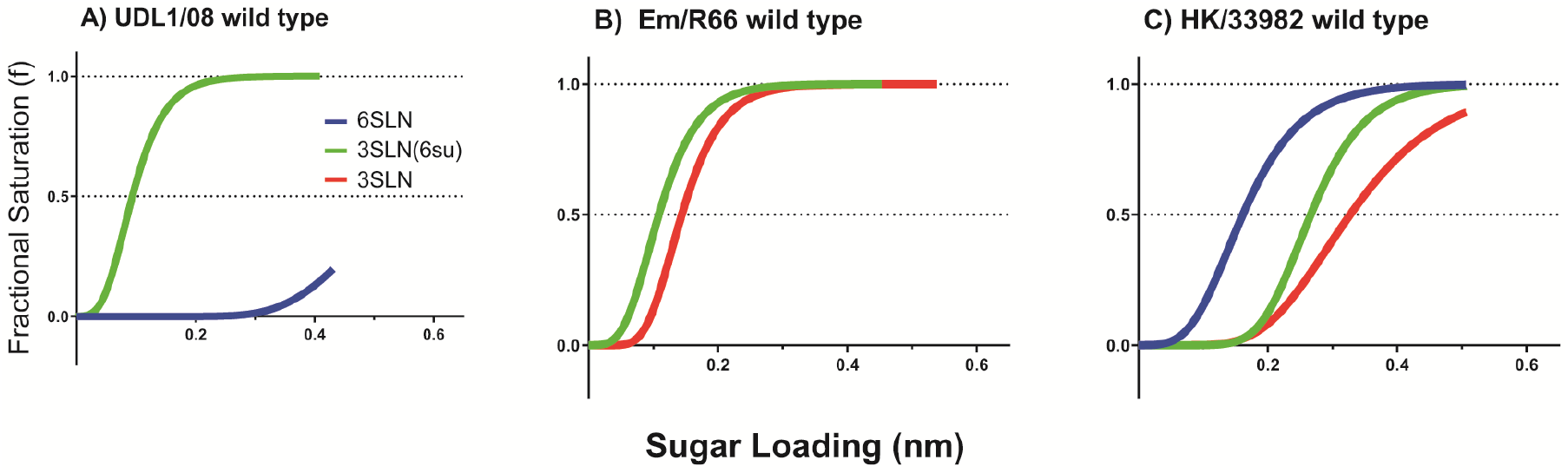
Varying receptor binding profiles of naturally occurring viruses. Biolayer interferometry was used to determine the receptor binding profiles of H9N2 viruses UDL1/08 (Panel A), Em/R66 (B), and HK/33982 (C). Binding was measured to three receptor analogues representative of: 1) an avian-like receptor (α2,3-sialyllactosamine or 3SLN, red lines), 2) a sulphated avian-like receptor (Neu5Ac α2,3 Gal β1-4(6-HSO3)GlcNAc or 3SLN(6su), green lines), and 3) a human-like receptor (α2,6-sialyllactosamine or 6SLN, blue lines). Where no binding was observed, the lines are omitted.

### Molecular basis of preference for sulphated and non-sulphated avian receptors

We previously showed that contemporary H9N2 viruses generally have a strong preference for sulphated avian-like receptors, a property not shared with non-H9N2 avian influenza viruses. To explore the molecular basis of this preference, we investigated amino acid differences between UDL1/08 and Em/R66, which do and do not show this preference respectively. Generally, these viruses had few differences near the RBS, though they did differ at positions 190, 226, and 227. Mutants were tested with individual reciprocal substitutions at positions 190, 226, and 227 and combinations thereof.

Substituting all three variable RBS residues (190/226/227) led to an approximate switch of the receptor-binding phenotypes between the two viruses. Substituting these three residues increased UDL1/08 3SLN binding and eliminated its 6SLN binding and abolished Em/R66’s 3SLN binding while increasing its 6SLN binding (Figure 2, purple lines). In general, 190/226 reciprocal mutants expressed similar binding phenotypes to the triple mutants (Figure 2, brown lines), suggesting the L227I and I227L substitutions had only a modest influence on receptor-binding. The exception to this was that Em/R66 E190A/Q226L/L227I did not show the increased binding to 6SLN observed with Em/R66 E190A/Q226L (Figure 1E). However, in both cases the exact binding avidities to the three receptor analogues were not completely recapitulated; both triple mutants (UDL1/08 A190E/L226Q/I227L and Em/R66 E190A/Q226L/L227I) had reduced avidity compared to their wild-type parental viruses indicating that further substitutions are required to fully recover the receptor-binding profile of the donor viruses. Nonetheless, positions 190, 226, and to a lesser extent, 227 appeared to be the primary determinants of variation in receptor-binding preference phenotypes of UDL1/08 and Em/R66s (Table 1).

**Figure 2 -.**
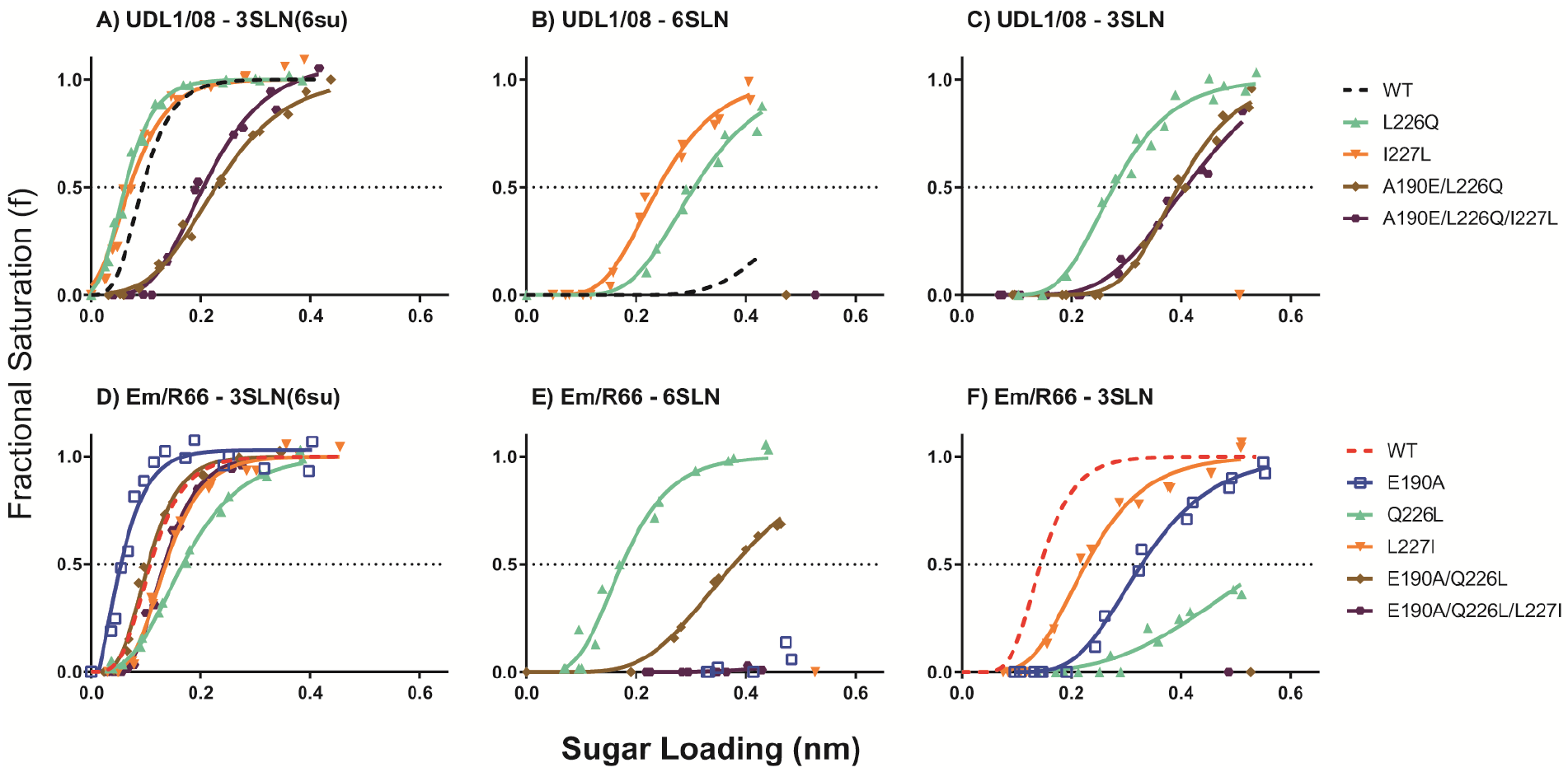
Receptor binding profiles of reciprocal UDL1/08-Em/R66 mutants. Biolayer interferometry was used to determine the receptor binding profiles mutant UDL1/08 (Panels A-C, UDL1/08 A190E could not be rescued) and Em/R66 viruses (panels D-F). Binding was measured to 3 receptor analogues: Neu5Ac α2,3 Gal β1-4(6-HSO3)GlcNAc (3SLN(6su), Panels A,D), α2,6-sialyllactosamine (6SLN, Panels B,E) and α2,3-sialyllactosamine (3SLN, Panels C,F). Dashed black and red lines show wild-type UDL1/08 and Em/R66 binding, respectively.

**Table 1.** Haemagglutinin amino acid differences modulating preference for sulphated and non-sulphated avian receptor. ^a^ ‘=’ indicates <2 fold difference, ‘+/-’ indicates 2-10 fold increase/decrease, ‘++/--’ indicates 10-100-fold increase/decrease, ‘+++/’ indicates >100-fold increase/decrease in binding relative to the wild type virus. ‘null’ indicates no difference was able to be seen because no binding to this analogue was detected with or without the substitution. ‘DNR’ indicates the virus was unable to be rescued. ‘/’ indicates a mutant was not made in that virus background.

| Residue (H3 no) | Residue (H9 no) | WT Virus | Substitution | Receptor |  |
| --- | --- | --- | --- | --- | --- |
|  |  |  |  | 3SLN(6su) | 3SLN |
| 190 | 180 | UDL1/08 | A->E | DNR | DNR |
|  |  | Em/R66 | E->A | ++ | -- |
| 190/226 | 180/216 | UDL1/08 | A->E, L->Q | -- | + |
|  |  | Em/R66 | E->A, Q->L | = | --- |
| 190/226/227 | 180/216/217 | UDL1/08 | A->E, L->Q, I->L | -- | + |
|  |  | Em/R66 | E->A, Q->L, L->I | - | --- |

Further investigating the contribution of individual RBS residues, position 190 exerted a strong influence on the preference for sulphated or non-sulphated avian-like receptors; viruses with A190 showed increased binding to 3SLN(6su) and decreased binding to non-sulphated 3SLN analogues while those with E190 showed the opposite (Figure 2A,C,D,F). This is potentially due to a charge repulsion between the negatively charged side-chain of glutamate and the sulphate group of sulphated-3SLN. Additionally, viruses containing A190 generally retained or had enhanced human-like 6SLN binding as seen in UDL1/08 wild-type and the mutants Em/R66 E190A and Em/R66 E190A/Q226L (Figure 2B, E). This was also exemplified in the differences in binding between UDL1/08 mutants: A190E/L226Q led to a loss of 6SLN binding while L226Q alone did not (Figure 1B – mint and brown lines).

At position 226, we determined that substitutions were exerting an effect on receptor-binding avidity, consistent with the results of our previous work with red blood cell based avidity assays [25]. Viruses with Q226 bound with higher avidity, compared with L226, to each of the three analogues tested (Figure 2A-F). Additionally, Q226 favoured 3SLN binding as can be seen by the mutant UDL1/08 L226Q and the difference in avidity between Em/R66 E190A/Q226L and E190A alone (Figure 2C, F – blue and brown lines). In the background of Em/R66, Q226L showed a large increase in 6SLN binding (Figure 2E – mint line), however a matching reduction in 6SLN binding by UDL1/08 L226Q was not seen (Figure 2B), indicating this is probably dependent on the context of the other amino acids in the H9 HA RBS as we had previously predicted [21].

Finally, substitutions at position 227 were identified as playing a minor role in modulation of avidity, though did not change receptor preference or regulate complete gain or loss of binding to any analogue. Relative to the associated parental virus, UDL1/08 I227L showed higher avidity to all receptors while Em/R66 L227I showed lower avidity (Figure 2, orange lines), consistent with previous inferences made from indirect measurements of avidity [25].

### Molecular basis of preference for the human receptor

To investigate the molecular basis of human-like receptor-binding as seen in some H9N2 viruses, a further reciprocal library was generated between UDL1/08 and HK/33982, a virus isolated from a human case of H9N2. UDL1/08 naturally binds to sulphated 3SLN with slight binding to 6SLN, while HK/33982 binds strongly to the human-like receptor 6SLN with moderate binding to both sulphated and non-sulphated 3SLN (Figure 1A, C). The amino acids at the three residues shown above to largely determine preference for sulphated vs non-sulphated (190, 226, and 227) also vary between these two viruses. In addition to these three positions, reciprocal mutations were introduced at positions 188 and 193 (178 and 183 in H9 numbering) on the basis that these residues are located next to the binding site and vary between UDL1/08 and HK/33982.

Introducing reciprocal mutations at the three residues (190/226/227) shown above to modulate preference for the sulphated vs non-sulphated avian receptor also influenced receptor preference in this case. Introducing the substitutions A190D, L226Q and I227Q from HK/33982 into UDL1/08 led to a loss of most of its 3SLN(6su) binding and a slight gain in 6SLN binding (Figure 2A, B, purple lines), however this binding phenotype did not resemble that of HK/33982. This suggests additional substitutions are required for a full gain of the human-adapted receptor-binding profile. The reciprocal triple mutant was unable to be recovered in the HK/33982 background, indicating additional residues must play a role in stabilising residues 190, 226 and 227 between these two viruses (Table S1).

Introduction of reciprocal substitutions at residues 190 and 227 together indicated that these positions likely play an important role in modulation of preference between the sulphated avian and human-like receptors. UDL1/08 with A190D/I227Q showed slightly decreased binding to 3SLN(6su) and appreciably increased 6SLN binding (Figure 3A, B – grey lines). HK/33982 D190A/Q227I had reduced receptor-binding to 6SLN and increased binding to sulphated 3SLN, creating an approximation of the receptor-binding preference of wild-type UDL1/08 (Figure 3D, E – grey lines). From this mutant library it is clear that switches at position 226 create an incompatibility in both virus backgrounds, with several viruses losing nearly all receptor-binding (UDL1/08 A190D/L226Q and UDL1/08 A190D/L226Q/I227Q) or infectious virus unable to be rescued entirely (HK/33982 D190A/Q226L and HK/33982 D190A/Q226L/Q227I) (Table S1).

**Figure 3.**
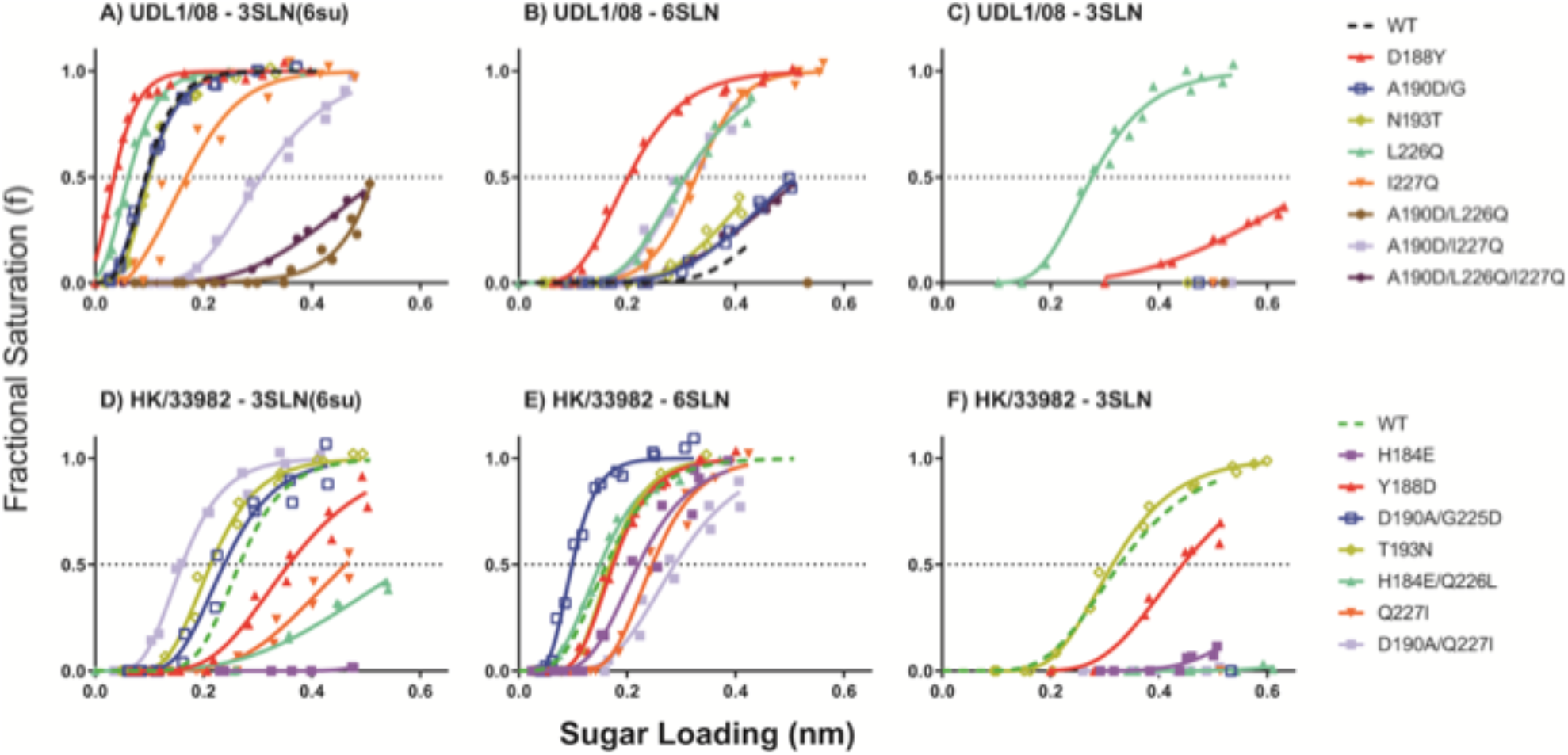
Receptor binding profiles of reciprocal UDL1/08-HK/33982 mutants. Biolayer interferometry was used to determine the receptor binding profiles mutant UDL1/08 (Panels A-C) and HK/33982 viruses (panels D-F). Binding was measured to 3 receptor analogues: Neu5Ac α2,3 Gal β1-4(6-HSO3)GlcNAc (3SLN(6su), Panels A,D), α2,6-sialyllactosamine (6SLN, Panels B,E) and α2,3-sialyllactosamine (3SLN, Panels C,F). Dashed black and green lines show wild-type UDL1/08 and HK/33982 binding, respectively.

The effects of single amino acid substitutions made between UDL1/08 and HK/33982 were also evaluated. Amino acid substitutions at residue 190 in the single reciprocal mutants of UDL1/08 and HK/33982 showed potential incompatibilities. UDL1/08 A190D showed a mixed population also including D190G while HK/33982 D190A gained the additional substitution G225D, as previously described [25]; HK/33982 with the G225D substitution alone was also unable to produce infectious virus (Table S1). It is, therefore, hard to draw conclusions from these viruses upon the influence of these mutations in isolation. HK/33982 D190A/G225D did appear to increase 6SLN binding and 3SLN(6su) binding while decreasing 3SLN binding (similar to Em/R66 E190A) (Figure 3D-F – blue lines). UDL1/08 A190D/G showed no difference in binding compared to wild-type UDL1/08 (Figure 2A-C – blue lines), however it is difficult to interpret these data, given the mixed population.

**Table 2.**
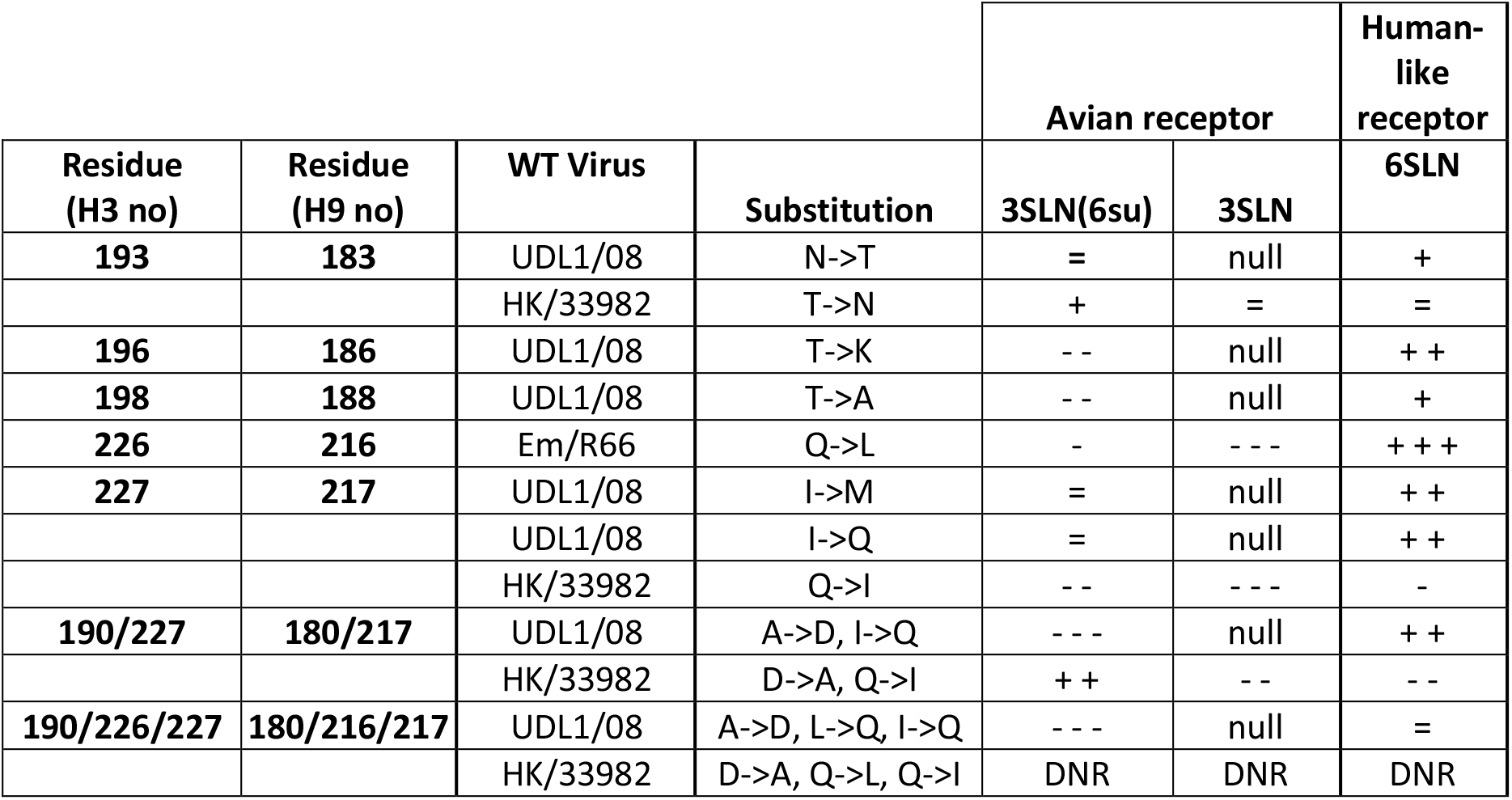
Haemagglutinin amino acid differences modulating preference for avian and human-like receptors. ^a^ ‘=’ indicates <2 fold difference, ‘+/-’ indicates 2-10 fold increase/decrease, ‘++/--’ indicates 10-100-fold increase/decrease, ‘+++/---’ indicates >100-fold increase/decrease in binding relative to the wild type virus. ‘null’ indicates no difference was able to be seen because no binding to this analogue was detected with or without the substitution. ‘DNR’ indicates the virus was unable to be rescued. ‘/’ indicates a mutant was not made in that virus background.

|  |  |  |  | Avian receptor |  | Human-like receptor |
| --- | --- | --- | --- | --- | --- | --- |
| Residue (H3 no) | Residue (H9 no) | WT Virus | Substitution | 3SLN(6su) | 3SLN | 6SLN |
| 193 | 183 | UDL1/08 | N->T | = | null | + |
|  |  | HK/33982 | T->N | + | = | = |
| 196 | 186 | UDL1/08 | T->K | -- | null | ++ |
| 198 | 188 | UDL1/08 | T->A | -- | null | + |
| 226 | 216 | Em/R66 | Q->L | - | --- | +++ |
| 227 | 217 | UDL1/08 | I->M | = | null | ++ |
|  |  | UDL1/08 | I->Q | = | null | ++ |
|  |  | HK/33982 | Q->I | -- | --- | - |
| 190/227 | 180/217 | UDL1/08 | A->D, I->Q | --- | null | ++ |
|  |  | HK/33982 | D->A, Q->I | ++ | -- | -- |
| 190/226/227 | 180/216/217 | UDL1/08 | A->D, L->Q, I->Q | --- | null | = |
|  |  | HK/33982 | D->A, Q->L, Q->I | DNR | DNR | DNR |

Similar to the reciprocal mutants of UDL1/08 and Em/R66, amino acid substitutions at residue 226 between UDL1/08 and HK/33982 displayed a clear avidity effect, with viruses possessing 226L appearing to show higher avidity compared with 226Q. HK/33982 with Q226L gained the additional compensatory substitution H184E (Table S1); HK/33982 with H184E alone showed a large reduction in avidity to all receptor analogues, whereas HK/33982 with H184E/Q226L increased avidity to both 6SLN and 3SLN(6su) relative to H184E alone, suggesting Q226L increased binding avidity (Figure 3D-F – purple and mint lines). The mutant HK/33982 H184E/Q226L also showed a strong preference for 6SLN.

At position 227, the introduction of I227Q in the UDL1/08 background resulted in a large increase in 6SLN binding, a drop in binding to UDL1/08’s preferred receptor 3SLN(6su), and did not alter binding to 3SLN (Figure 3A-C, orange lines). The reciprocal substitution, HK/33982 Q227I, showed a general avidity effect with lower binding to all analogues (Figure 3D-F – orange line), however when introduced alongside D190A, the impact of Q227I in the HK/33982 D190A/Q227I double mutant facilitated reduced 6SLN binding but slightly increased 3SLN(6su), relative to the parental wild-type and the D190A(+G225D) mutant (Figure 3D-F – grey and orange lines).

Reciprocal substitutions at position 188 were found to influence receptor-binding while those at position 193 were not. These residues have not previously been described as affecting receptor-binding in H9 HA, though they are at the edge of the RBS and 193 has been described as playing a vital role in the modulation of binding of sulphated 3SLN by H5 and H7 HA [26, 27]. Viruses with Y188 showed higher avidity relative to viruses with D188, regardless of the virus background (Figure 3 – red lines). Additionally, HK/33982 Y188D appeared to have minor effects on specific receptor analogue preference; this substitution reduced binding for the 3SLN analogues to a greater extent than the 6SLN analogue. Amino acid swaps at position 193 showed a very minor, non-reciprocal receptor preference effect (Figure 3), consistent with our previous estimates that swaps of residue 193 between UDL1/08 and HK/33982 would only have a minimal receptor avidity effect [25].

### Molecular basis of variation in receptor-binding avidity

In addition to showing preference for different receptors, influenza viruses vary in receptor-binding avidity. In several previous studies we have inferred that escape mutants with the largest impact of polyclonal antisera binding may be driven by avidity effects [20, 25, 28]. To further investigate the genetic basis of variation in receptor-binding avidity we constructed a library of mutants in the UDL1/08 background and assessed their receptor-binding phenotypes. To complement this, we analysed a large dataset of haemagglutination inhibition (HI) titres and HA sequences from natural H9N2 viruses to identify amino acid variants correlating with avidity effects apparent in the measured HI titres.

Testing of UDL1/08 mutants showed several to exhibit a general decrease in avidity to the tested analogues including K137I, F147L, K157T, N158D, N193D, and G225D (Figure 4A, B). Only a single substitution, T189N, showed a negligible effect on receptor-binding (Figure 4C, D - mint lines). Two substitutions, T135K and T192R, appeared to facilitate a general increase in avidity (i.e. increased binding to all receptor analogues tested) in a similar manner to UDL1/08 I227L and D188Y from the reciprocal mutant libraries (Figure 4C, D, 2A-C and 3A-C). R82G showed a reduction in 3SLN(6su) binding while not having an effect on 6SLN binding (Figure 4E,F – blue lines). Finally, one group of mutants showed changes in receptor-binding preference with increases in ‘human-like’ 6SLN binding relative to ‘avian-like’ 3SLN(6su): I227M showed a large increase in 6SLN binding without changing sulphated 3SLN binding (orange lines); G159K showed a modest increase in avidity to sulphated 3SLN but a much larger relative increase in 6SLN binding (purple lines); T196I, T196K, and T198A all decreased sulphated 3SLN binding while increasing 6SLN binding to varying degrees (Figure 4E,F – light red, dark red and light brown lines). This last group of mutations represent single amino acid changes that could act as markers for viruses with greater zoonotic potential, along with the previously described Q226L (in the background of Em/R66 and HK/33982), and I227Q in a UDL1/08-like background (Figure 2E, 3B,E).

**Figure 4.**
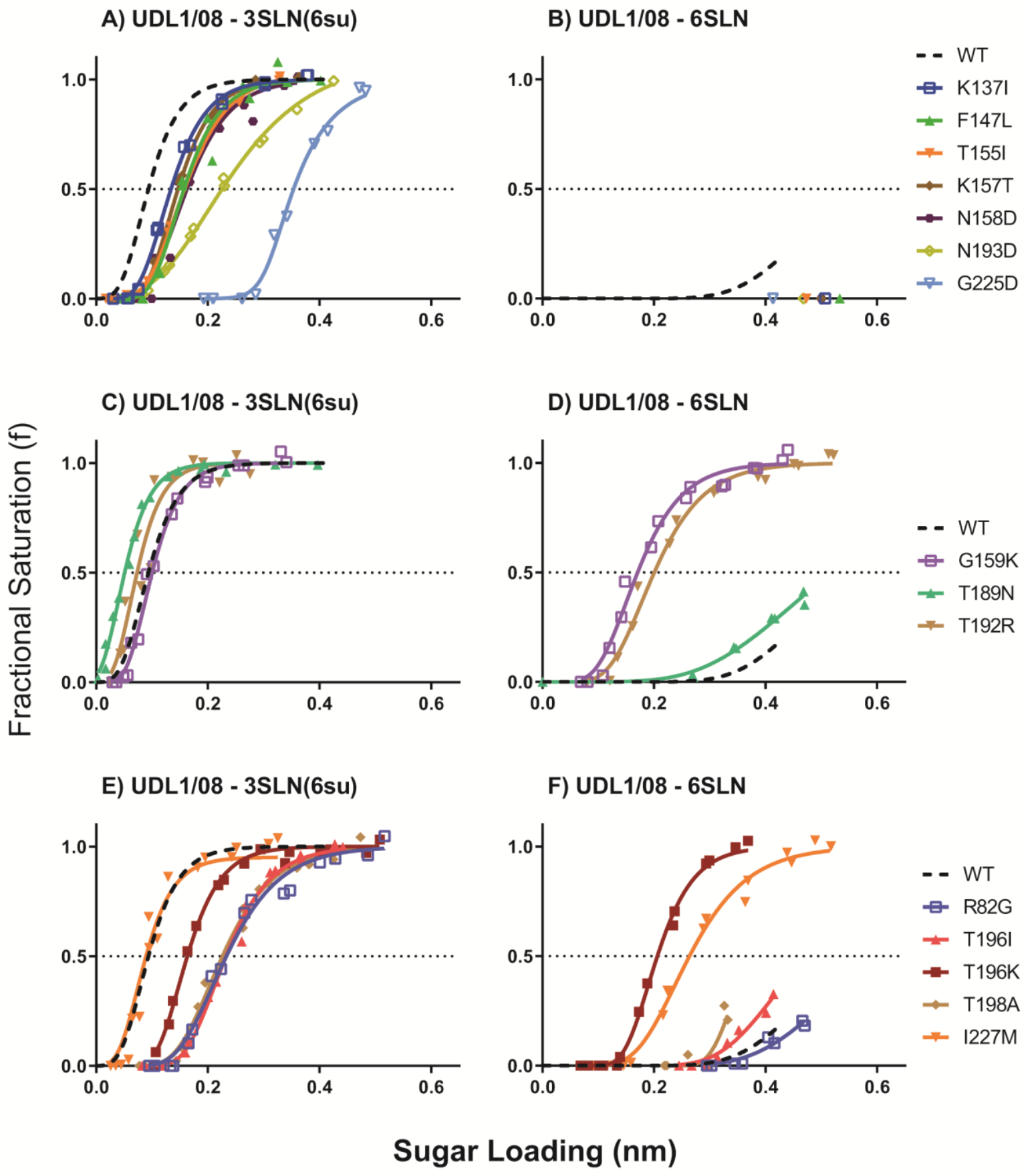
Receptor binding modulation by non-reciprocal mutations in the background of UDL1/08. Biolayer interferometry was used to determine the receptor binding profiles of each mutant virus. Binding was measured to 3 receptor analogues, α2,6-sialyllactosamine (6SLN), α2,3-sialyllactosamine (3SLN) and Neu5Ac α2,3 Gal β1-4(6-HSO3)GlcNAc (3SLN(6su)). No viruses had any detectable 3SLN binding. Dashed black lines show wild-type UDL1/08 binding. Panels indicate binding by mutants that show an avidity reduction (A,B), an increase in avidity (C,D) and changes in receptor binding preference (E,F). No mutants had any detectable binding to 3SLN.

In general, amino acid replacements that increased positive charge in the HA head domain tended to cause an increase in receptor-binding avidity while the opposite was true for substitutions that increased negative charge. In Figure 5A, the impact of substitutions introduced to the UDL1/08 backbone on avidity for its preferred receptor, 3SLN(6su), is plotted by the change in net charge of the HA head domain. The proximity of the residues at which substitutions were introduced to the RBS is shown in Figure 5B.

**Figure 5 -.**
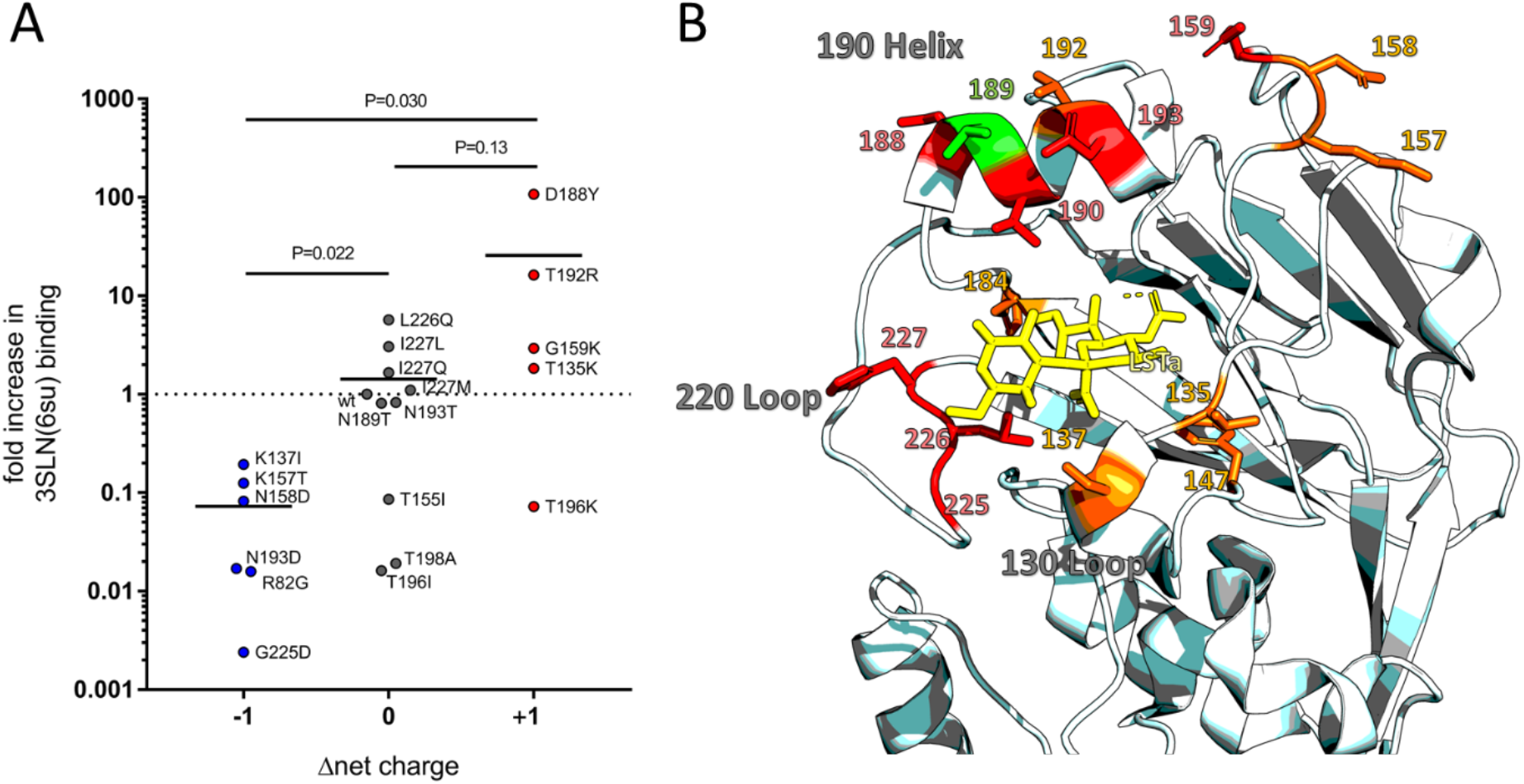
Mapping of residues tested for receptor binding changes and correlation between charge and 3SLN(6su) binding. A) The fold-increase in binding to 3SLN(6su) is plotted along with the introduced change in net charge difference. Wild type UDL1/08 binding to 3SLN(6su) is set to 1 with relative binding calculated as described previously [21]. Annotated P values calculated using a non-parametric Mann-Whitney U test – lines indicate mean fold increase. B) Residues which effect receptor binding preference (i.e. effect binding to different analogues in different ways) shown in red, residues that affect receptor binding avidity across analogues shown in orange. Residue that when mutated have no effect on receptor binding shown in green. LSTa, shown in yellow, is an α2,3-linked (avian-like) receptor analogue. H3 HA numbering used throughout. Figure made using structure PDBID:1JSH [41], made using PyMol [42].

To investigate variation in avidity among natural H9N2 viruses, we modelled variation in HI titres for a dataset of large number of viruses covering all major H9 lineages. In addition to measuring antigenic similarity of influenza viruses, HI titres are influenced by varying avidity. Viruses that bind receptors on the surface of red blood cells used in the assay with higher avidity require more antibodies to inhibit haemagglutination manifesting as a tendency towards lower HI titres regardless of antigenic relationships to the antisera used, and vice-versa. To identify amino acid variants correlating with such variation in titres, we adapted a model we previously developed to explore antigenic variation in both human and avian influenza viruses, identifying several substitutions predicted to cause antigenic variation and validating these using mutagenesis [25, 29].

Variation in HI titres resulting both from antigenic differences and from variation in virus avidity was mapped to branches of the H9 HA phylogenetic tree as previously described [25]. To explore the genetic basis of variation caused by differences in avidity, phylogenetic terms representing branches leading to clades of viruses with systematically higher or lower titres were replaced with terms representing amino acid identity in the test virus at each variable HA position in turn. Under a forward selection procedure, terms representing positions 190, 196 and 198 were selected (180, 186 and 188 in H9 numbering, respectively). Each of these positions are in the 190-helix, proximal to the RBS. Position 190 has already been shown to play a role in receptor-binding in this and other studies [18, 20], and we see the effect of 196 and 198 on receptor-binding preference in this study (Figure 4E,F), further validating this modelling approach (Table 3).

**Table 3.** Amino acid residues predicted to explain variation in HI titres as a result differences in receptor-binding avidity. Effect on 3SLN(6su) binding is shown relative to most common amino acid in meta-analysis dataset: 190A, 196T, and 198T.

| Residue (H3 no) | Residue (H9 no) | Distinct amino acid residues<sup>b</sup> | Number of HI titres | Effect on 3SLN(6su) binding |
| --- | --- | --- | --- | --- |
| 190 | 180 | A | 1,145 |  |
|  |  | T | 309 | 50x increase |
|  |  | V | 494 | 1400x increase |
| 196 | 186 | T | 1,912 |  |
|  |  | I | 331 | 41x decrease |
|  |  | K | 296 | 7x decrease |
| 198 | 188 | T | 1,913 |  |
|  |  | A | 351 | 41x decrease |

### Sequence-based prediction of receptor-binding preference of H9N2 viruses

Finally, we extrapolated the results of this study to predict the receptor-binding preferences of circulating H9N2 viruses on the basis of amino acid identity at positions 190, 226 and 227. Viruses were predicted as possessing one of three receptor-binding phenotypes (Figure 6): 1) a strong preference for sulphated avian-like receptors, the canonical chicken-adapted H9N2 virus phenotype (shown in green and represented by UDL1/08), 2) a receptor-binding phenotype more similar to chicken-adapted H5Nx or H7N1 viruses and binds both sulphated and non-sulphated avian-like receptors (shown in red and represented by Em/R66), or 3) an preference for the human-like receptor with concurrent binding to avian receptors which we hypothesise may be an adaptation to minor poultry (shown in blue and represented by HK/33982). Viruses were predicted to exhibit preference for the sulphated avian-like receptor if at positions 190-226-227 they possessed either A, T, or V at 190, L at 226 and either I, L, M, or Q at 227, or if they possessed the motifs A-Q-I, A-Q-T, or I-Q-F. Viruses possessing either E-Q-L or E-Q-Q were predicted to bind both sulphated and non-sulphated avian-like receptors. Viruses possessing A, D, T, or V at 190 and Q-Q at 226-227 were predicted to show preference for the human-like receptor, as were viruses possessing the motif E-L-Q. Predictions for motifs without direct evidence but made based on combinations of other motifs results are shown in lighter shades to indicate reduced confidence (details in Materials and Methods), while predicted receptor preference was considered unknown for viruses with incomplete sequence information at positions 190, 226 and 227 or alternative motifs. The substitutions at positions 196 and 198 that increased relative 6SLN binding were not included at this stage as it is not known with which other substitutions must occur to elevate 6SLN binding above binding for the avian receptor.

**Figure 6.**
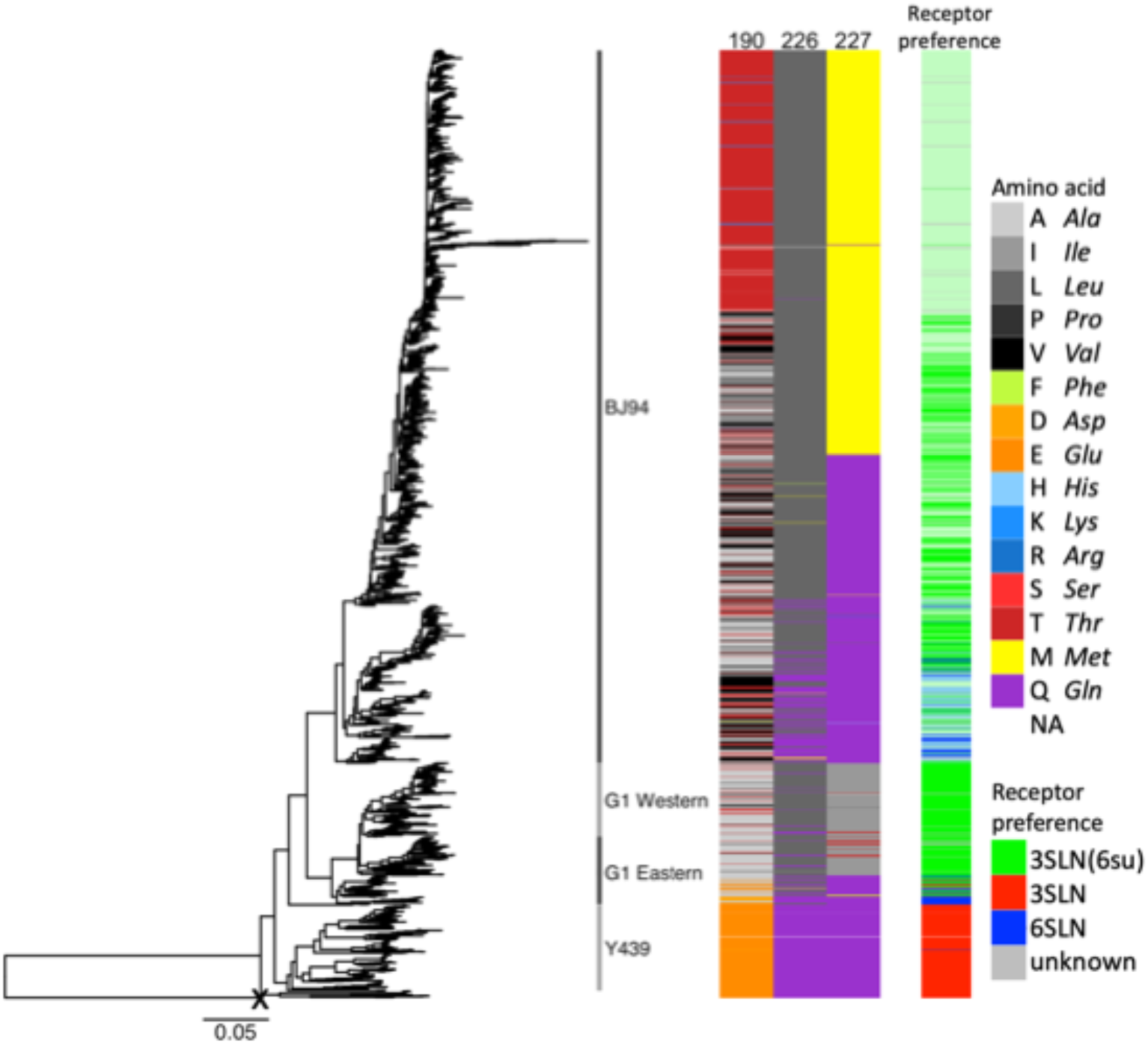
Sequence variation and predicted receptor preference across H9N2 phylogeny. HA phylogeny for 2,440 H9N2 viruses generated using and rooted with a sequence from H9-like bat influenza virus (starred). Amino acid identity at positions 190, 226 and 217 is shown by colour, grouped by side-chain property, according to the legend. Predicted receptor preference for α2,6-sialyllactosamine (6SLN), α2,3-sialyllactosamine (3SLN) and Neu5Ac α2,3 Gal ϐ1-4(6-HSO3)GlcNAc (3SLN(6su)) is shown in blue, red and green respectively, with lighter shades indicating reduced confidence. Predicted receptor preference is based on amino acid identity at positions 190, 226 and 227 based on extrapolation of the amino acid sequence of viruses tested using BLI and the results of mutagenesis experiments.

Amino acid sequence at positions 190, 226, and 227 and the resulting prediction of receptor preference is shown across a phylogeny constructed using all available H9N2 HA sequences in Figure 6. Almost all viruses in Y439-like lineage, prevalent in wild birds and poultry in Korea, as well as a few viruses in the G1-Eastern sub-lineage are predicted to show an Em/R66-like preference for any avian-like receptor. The vast majority of viruses in the chicken-adapted BJ94 and G1 lineages are predicted to maintain a canonical H9N2 sulphated avian receptor preference. A significant number of viruses belonging to the G1-Eastern sub-lineage, prevalent in minor poultry in China, are predicted to show a preference for human-like receptors, as are a number of viruses interspersed within a clade of viruses belonging to the BJ94 lineage.

## Discussion

Receptor-binding is an important determinant of host specificity and modification of receptor binding properties is often a critical step in cross-species virus transmission. In this study we have comprehensively investigated residues in and around the receptor binding site of HA from H9N2 viruses for their ability to influence receptor preference and avidity. We have shown that different combinations of the residues 190, 226 and 227 account for much, but not all, of the variability in H9N2 receptor binding between viruses representative of the major binding phenotypes. Furthermore, we described several other residues that have strong influences on H9N2 receptor binding preference including positions 159, 188, 193, 196 and 198. We propose a model whereby residues in the influenza HA1 that don’t directly coordinate receptor binding can play a delocalised avidity role through modulating the charge of the head domain with increases in positive charge giving a non-specific increase in avidity and vice versa. We hypothesise this effect may be exaggerated for H9N2 viruses where sulphated, siaylated glycans appear to be the preferred receptor owing to the greater amount of negative charge in these receptors (compared to non-sulphated, siaylated glycans). Finally, we have applied the results of this study to try and predict the receptor binding phenotypes of different circulating H9N2 viruses as a way of predicting strains that may have a heightened zoonotic potential.

Several of the residues identified in this study have been previously described as directly or indirectly affecting H9 receptor binding, including position 155, 190, 225, 226 and 227 [13, 17–20, 30, 31]. In previous studies, generally only one or two of these residues are measured in isolation for their receptor binding effect or are tested in a non- or only semi-quantitative manner. Here we perform a comprehensive, quantitative analysis often testing changes at residues in multiple different virus backbones and in multiple combinations to investigate strain-specific and compensatory effects. For example, residue 226 has been shown several times in an older G1-Eastern lineage virus to increase binding to human-like receptors [17, 32], with a similar effect when we investigate a contemporary virus of the same lineage (i.e. HK/33982), however in a G1-Western backbone (i.e. UDL1/08) we find a completely different effect whereby there is a general avidity increase and an overall increase in 6SLN-binding when a L226Q substitution is incorporated. This highlights the importance of the overall context of receptor binding residues when looking at influenza receptor binding mutants.

In our previous study we predicted that one of the groups of substitutions that had the largest effect on immune escape in H9N2 viruses were substitutions that affect avidity, as has been predicted for human influenza viruses [25, 33, 34]. Here we test a wide variety of escape mutants that were previously shown to robustly modulate polyclonal antisera binding and show a large number of them also modulate receptor binding avidity. This suggests that many escape mutations that have a large influence on antigenicity may be exerting this effect through a receptor binding avidity effect as has been shown for human influenza viruses [33, 34]. We also performed an analysis of matched genetic and antigenic data for 330 H9N2 viruses, covering each of the major H9 lineages [25]. Using a modified version of a model used to identify antigenic determinants, we predicted that different residues at positions 190, 196 and 198 would explain variation in HI titres as a result of differing avidity, as they tended to be associated with lower or higher HI titres irrespective of antigenic relationships between the viruses and antisera being compared. Each of these three positions are confirmed in this study to play an important role in regulation of receptor-binding avidity. In addition to the important role for 190 in receptor preference, the substitutions T196I, T196K and T198A all showed relative increases in human-like receptor binding. These results further demonstrate there is a strong case for using integrated modelling approaches to reanalyse large data sets and predict residues that affect receptor binding and antigenic phenotypes.

We present evidence suggesting that residues in the influenza HA1 that don’t directly coordinate receptor binding play a delocalised avidity role through modulating the charge of the head domain. For mutations introduced in the UDL1/08 background and measured in binding to its preferred receptor, 3SLN(6su), we found a significant trend in avidity change between substitutions that introduced a negative charge and those that introduced a positive charge. In general, substitutions observed to increase avidity tended to increase the net positive charge around the RBS (e.g. T135K, G159K, T192R, D188Y, Em/R66 E190A, Em/R66 Q226L/E190A) whilst avidity decreasing mutants usually decreased the net positive charge around the RBS (e.g. R82G, K137I, K157T, N158D, N193D, G225D, HK/33982 Y188D, HK/33982 H184E). This effect is likely due to non-specific charge interactions with the negatively charge sialic acid and we hypothesise may be more pronounced for H9N2 viruses as the negative charge of sulphated, siaylated glycans, which appear to be their preferred receptor, is greater due to the negatively charged sulphate group.

An important contribution of this study is the identification of several substitutions that, in one or more of our virus backbones, resulted in viruses with increased or *de novo* human-like 6SLN binding. These residues will be useful for future surveillance efforts to identify newly sequenced viruses with elevated zoonotic potential. These particular substitutions include the previously well characterised Q226L substitution in HK/33982 or Em/R66-like backbones (but not the contemporary G1-Western UDL1/08-like backbone) as well as the newly characterised R82G, T135K, G159K, T196I, T196K, T198A, I227M and I227Q substitutions in the UDL1/08-like backbone. Several of these mutations are already commonly found in the field, further suggesting that H9N2 virus variants naturally circulate with a likely heightened zoonotic potential.

In conclusion, we have quantified the impact of single and multiple amino acid substitutions on receptor binding phenotypes in the context of several H9N2 viruses with varying receptor binding preferences, identifying seven novel mutations that increase binding to the human-like receptor. We further highlight the importance of mutations that impact receptor binding avidity. Avidity modulation has a dramatic impact on antigenicity and an equally important role in receptor binding phenotype, thus viruses that gain avidity enhancing mutations may present multiple challenges in that both vaccine efficacy may be compromised, and zoonotic potential may be increased concurrently. As well as helping better understand the molecular basis of avian influenza receptor binding, the results generated here will help future surveillance efforts to identify viruses which may potentially have an augmented zoonotic potential and/or greater vaccine escape potential.

## Materials and Methods

### Ethics statement

Use of embryonated eggs in this study was carried out in strict accordance with European and United Kingdom Home Office regulations and the Animals (Scientific Procedures) Act 1986 Amendment Regulations, 2012. These studies were carried out under the United Kingdom Home Office approved project license numbers 30/2683 and 30/2952.

### Cell lines and eggs

HEK 293Ts and MDCKs were maintained in Dulbecco’s modified Eagle medium (DMEM), supplemented with 10% foetal calf serum, 37°C, 5% CO_2_. Viruses were propagated in 10-day-old embryonated eggs; allantoic fluid was harvest 48 hours post-inoculation.

### Viruses

Throughout this study recombinant viruses, generated by standard 8 plasmid influenza reverse genetics were used [35]. All viruses contained the named HA gene (whether wild type or mutant), the NA of A/chicken/Pakistan/UDL-01/2008 (UDL1/08) and the remaining genes from A/Puerto Rico/8/1934 (PR8), allowing for high viral titres from eggs. Mutant HA plasmids were generated by site directed mutagenesis. Viruses were attempted to be rescued a minimum of three independent times and left for 7 days post-co-culture before being determined to be un-rescuable.

### Virus sequencing

Viruses were sequenced to confirm no reversions or additional substitutions had occurred upon production and propagation. The HA1 region of HA was sequenced for each virus as previously described [36].

### Virus purification

Low speed centrifugation was initially used to remove large debris from virus containing egg allantoic fluid. Virus particles were next pelleted by ultracentrifugation at 27,000rpm for 2 hours. Virus pellets were subsequently homogenised by glass homogeniser, resuspended and purified with a 30-60% sucrose gradient. The visible band containing virus was then isolated, diluted into PBS and centrifuged for another 2 hours at 27,000rpm. The final virus pellet was then resuspended in PBS, 0.01% azide. Concentration of purified viruses was determined using a nucleoprotein ELISA as described previously [37].

### Biolayer interferometry

Purified virus binding to different sialylated receptor analogues was tested using an Octet RED bio-layer interferometer (Pall ForteBio) as described previously [21]. Receptor analogues contained 30kDa polyacrylamide backbones conjugated to 20 mol % trisaccharides and 5 mol % biotin (Lectinity Holdings). The three analogues used in this study were α2,6-sialyllactosamine (6SLN), α-2,3-sialyllactosamine (3SLN) and Neu5Ac α-2,3Galβ1-4(6-HSO3)GlcNAc(3SLN(6su)). Sialoglycopolyemers were bound onto streptavidin coated biosensors (Pall ForteBio) at ranges of concentrations from 0.01-0.55μg/ml in 10mM HEPES, pH 7.4, 150mM NaCl, 3mM EDTA, 0.005% Tween-20 (HBS-EP). Virus was diluted to a concentration of 100pM in HBS-EP, 10μM oseltamavir carboxylate (Roche), 10μM zanamivir (GSK). Virus association to the bound sialoglcopolymers was measured at 20°C for 30 minutes. Virus binding curves were normalised to fractional saturation and plotted as a function of sugar loading. Relative dissociation constants were calculated as described previously [21, 38].

### Modelling of potential receptor binding residues

To identify amino acid positions where substitutions correlated with differences in receptor-binding avidity apparent in HI titres, a modelling approach previously used to identify substitutions causing antigenic differences among influenza viruses was adapted [25, 29, 39, 40]. Following the methodology described in the aforementioned studies, branches of the HA phylogenetic tree correlated with variation in HI titres when the branch 1) separated test virus and antisera, 2) descended the test virus, or 3) descended the virus used to generate antisera. These phylogenetic terms are interpreted as being associated with changes in 1) antigenicity, 2) receptor-binding avidity, and 3) immunogenicity, respectively. Internal branches of the phylogeny leading to clades of two or more viruses associated with systematically higher or lower titres (numbered 2 in previous sentences) and containing at least one virus also used as an antisera strain were removed, effectively dropping any terms from the model that explained variation in HI associated with differences in virus avidity. In their place, terms representing amino acid identity in the assayed virus at each variable HA position were tested. At each position, these terms allowed for titres to vary according to which amino acid residue the virus possessed to account for potential differences in contributions to avidity. These position-specific terms were added to the model under a forward selection procedure until the addition of further terms ceased to improve the model, as assessed by likelihood ratio test (p < 0.05) with a Holm-Bonferroni correction for multiple testing. At selected positions, effect sizes were estimated for each alternative amino acid relative to the amino acid found most commonly in the dataset.

### Prediction of receptor binding profiles from sequence

All available HA sequences from H9 viruses were downloaded from GIASID. A phylogenetic tree was generated from aligned nucleotide sequences using MEGA. Receptor-binding profiles were predicted across the phylogeny according to amino acid identify at positions 190, 226, and 227 on the basis of the BLI results derived during this study. Viruses were predicted as possessing either: 1) a strong preference for sulphated avian-like receptors, 2) a receptor binding phenotype more similar to chicken-adapted H5Nx or H7N1 viruses that bind both sulphated and non-sulphated avian-like receptors or 3) a preference for the human-like receptor with concurrent binding to avian receptors. Viruses were predicted to exhibit preference for the sulphated avian-like receptor if at positions 190-226-227 they possessed the motifs A-L-I, A-L-L, A-L-M, A-L-Q, A-Q-I, A-Q-T, I-Q-F, T-L-I, V-L-I (or with reduced confidence T-L-L, T-L-M, T-L-Q, V-L-L, V-L-M, or V-L-Q), for any avian-like receptor if they possessed the motifs E-Q-L or E-Q-Q, and for the human-like receptor if they possessed A-Q-Q, D-Q-Q, E-L-Q (or with reduced confidence T-Q-Q or V-Q-Q). Predictions made with reduced confidence indicate that we have not tested the exact combination of amino acids but that the prediction is consistent with other combinations barring unforeseen interactions between sites. For viruses with incomplete sequence information at positions 190, 226 and 227 or alternative motifs, no prediction was made.

## Acknowledgements

The authors would like to thank Dr Stephen Martin of The Francis Crick institute for use of his software for analysing the bio-layer interferometry results and Dr Jürgen Stech of the Friedrick Loeffler Institute for the haemagglutinin reverse genetics plasmid of A/chicken/Emirates/R66/2002 (H9N2).

## Author contributions

Conceptualisation - TP, WH. Data curation, Formal analysis, funding acquisition – RR & MI Investigation - TP, JES & WH. Methodology, Resources, Software- WH & RR Supervision – WB, RR & MI. Writing original draft – TP, JES & WH. Writing – review and editing – All

