## Supplementary Materials for "Genetic determinants of receptor-binding preference and zoonotic potential of H9N2 avian influenza viruses"

### Supplementary Figures

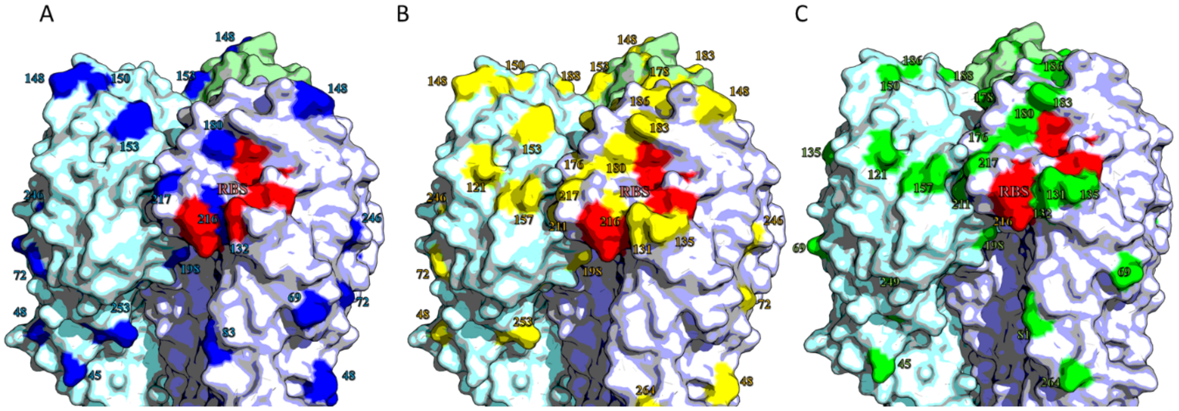


**Supplementary Figure 1. Amino acid differences in the head domain of H9 haemagglutinin between viruses used in this study.** A) Differences between UDL1/08 and Em/R66 shown in blue. B) Differences between UDL1/08 and Hk/33982 shown in yellow. C) Differences between Em/R66 and HK/33982 shown in green. Selected receptor binding residues shown in red. Figure made using structure PDBID:1JSH [41], made using PyMol [42].

### Supplementary Tables

Table S1. Attempted mutants that were not rescued or that were only rescued with compensatory changes or additional directed mutagenesis or showed a reversion.

| **Attempted mutant** | **Rescued?** | **Compensatory change** | **Additional substitution needed** | **Reverted** |
|---|---|---|---|---|
| UDL1/08 A190E | no | n/a | Yes, L226Q, I227I or I227Q | n/a |
| UDL1/08 A190D | yes | n/a | n/a | Yes, D190G (mixed, equal) |
| HK/33982 D190A | yes | Yes, G225D (total change[b]) | n/a | n/a |
| HK/33982 Q226L | yes | Yes, H184E (total change) | n/a | n/a |
| HK/33982 G225D | No | n/a | Yes, D190A | n/a |
| HK/33982 D190A/Q226L | No | n/a | n/a | n/a |
| HK/33982 D190A/Q226L/Q227I | No | n/a | n/a | n/a |
| HK/33982 Q227L | No | n/a | n/a | n/a |

[a]Indicates addition of potential glycosylation site.

[b]'Total change' indicates presence of singlet on.

[c]'Partial' indicates doublet with no change to consensus.

### Table S2. Summary of receptor binding changes seen in this study

| | | Effect of substitutions on WT virus[a] | | | | | | | | | | | |
|---|---|---|---|---|---|---|---|---|---|---|---|---|---|
| | | UDL1/08 | | | | Em/R66 | | | | HK/33982 | | | |
| Residue (H3 no) | Residue (H9 no) | Substitution | 3SLN(6su) | 6SLN | 3SLN | Substitution | 3SLN(6su) | 6SLN | 3SLN | Substitution | 3SLN(6su) | 6SLN | 3SLN |
| **single substitution** | | | | | | | | | | | | | |
| **82** | **74** | R->G | - - | = | ND | / | / | / | / | / | / | / | / |
| **135** | **129** | T->K | = | + | ND | / | / | / | / | / | / | / | / |
| **137** | **131** | K->I | - | - | ND | / | / | / | / | / | / | / | / |
| **147** | **137** | F->L | - - | - | ND | / | / | / | / | / | / | / | / |
| **155** | **145** | T->I | - - | - | ND | / | / | / | / | / | / | / | / |
| **157** | **147** | K->T | - | - | ND | / | / | / | / | / | / | / | / |
| **159** | **149** | G->K | + | +++ | ND | / | / | / | / | / | / | / | / |
| **158** | **148** | N->D | - - | - | ND | / | / | / | / | / | / | / | / |
| **184** | **174** | / | / | / | / | / | / | / | / | H->E | - - - | - | - - |
| **188** | **178** | D->Y | +++ | ++ | + | / | / | / | / | Y->D | - | = | - |
| **189** | **179** | N->T | = | = | ND | / | / | / | / | / | / | / | / |
| **190** | **180** | A->E | DNR | DNR | DNR | E->A | ++ | ND | - - | / | / | / | / |
| | | / | / | / | / | E->D | - - - | ND | - - - | D->E | - - | - - - | - |
| | | A->D/G | = | = | ND | / | / | / | / | / | / | / | / |
| | | A->V | +++ | ++ | + | / | / | / | / | / | / | / | / |
| | | A->T | ++ | ++ | ND | / | / | / | / | / | / | / | / |
| **192** | **182** | T->R | ++ | ++ | ND | / | / | / | / | / | / | / | / |
| **193** | **183** | N->D | - - | - | ND | / | / | / | / | / | / | / | / |
| | | N->T | = | + | ND | / | / | / | / | T->N | + | = | = |
| **196** | **186** | T->I | - - | = | ND | / | / | / | / | / | / | / | / |
| | | T->K | - - | ++ | ND | / | / | / | / | / | / | / | / |
| **198** | **188** | T->A | - - | + | ND | / | / | / | / | / | / | / | / |
| **225** | **215** | G->D | - - - | - | ND | / | / | / | / | G->D | DNR | DNR | DNR |
| **226** | **216** | L->Q | + | ++ | ++ | Q->L | - | +++ | - - - | / | / | / | / |
| **227** | **217** | I->L | + | ++ | ND | L->I | - | ND | - | / | / | / | / |
| | | I->Q | = | ++ | ND | / | / | / | / | Q->I | - - | - | - - - |
| | | I->M | = | ++ | ND | / | / | / | / | / | / | / | / |
| | | / | / | / | / | L->Q | - | ND | - | Q->L | DNR | DNR | DNR |
| **multiple substitutions** | | | | | | | | | | | | | |
| **184/226** | **174/216** | / | / | / | / | / | / | / | / | Q->L, H->E | - - | = | - - - |
| **190/225** | **180/215** | / | / | / | / | / | / | / | / | D->A, G->D | = | + | - - - |
| **190/226** | **180/216** | A->E, L->Q | - - | - | + | E->A, Q->L | = | + | - - - | / | / | / | / |
| **190/226** | **180/216** | | | | | / | / | / | / | D->A, Q->L | DNR | DNR | DNR |
| **190/227** | **180/217** | A->D, I->Q | - - - | ++ | ND | / | / | / | / | D->A, Q->I | ++ | - - | - - - |
| | | / | / | / | / | E->D, L->Q | - - | ND | - - | D->E, Q->L | - - - | - - - | - - - |
| **190/226/227** | **180/216/217** | A->E, L->Q, I->L | - - | - | + | E->A, Q->L, L->I | - | ND | - - - | / | / | / | / |
| | | A->D, L->Q, I->Q | - - - | = | ND | / | / | / | / | D->A, Q->L, Q->I | DNR | DNR | DNR |

[a] '=' indicates <2 fold difference, '+/-' indicates 2-10 fold increase/decrease, '++/- -' indicates 10-100-fold increase/decrease, '+++/- - -' indicates >100-fold increase/decrease in binding relative to the wild type virus. 'ND' indicates not difference was able to be seen because no binding to this analogue was detected with or without the substitution. 'DNR' indicates the virus was unable to be rescued. '/' indicates a mutant was not made in that virus background.
